# Deep Kernel Inversion: Rapid and Accurate Molecular Interaction Prediction for Drug Design

**DOI:** 10.1101/2024.11.19.624391

**Authors:** Seth A. Myers, Chad Miller, Kyler Lugo, Thi Trinh, Chang-wook Lee, Naveen Arunachalam, Hannah Chen, Denis Drygin, James Martineau

## Abstract

Computational drug design offers the opportunity to dramatically accelerate novel therapeutics for untreated diseases. Designing compounds with optimal efficacy and specificity, however, requires understanding and optimizing immense numbers of molecular interactions. While advances in predicting one-to-one molecular interactions continue, there has been limited progress in scaling one-to-many or many-to-many molecular interaction models. In this paper, we introduce a deep learning framework that embeds molecules into a high-dimensional vector space, which we have named Deep Kernel Inversion. In this framework, the dot product between vectors accurately predicts molecular interactions. This approach reduces the complexity of predicting an entire molecular interaction network from *O*(*n*^2^) to *O*(*n*), enabling new molecular design tasks previously inaccessible to computational approaches. In the case of human protein-protein interactions (PPI), we demonstrate a *100,000 fold* decrease in the computation required to map the full human PPI network. We also demonstrate best-in-class performance across multiple molecular interaction tasks with this approach. This work offers a new way forward in scaling accurate molecular interaction predictions with applications in mapping biological pathways, target discovery, drug design, and therapeutic development.

## 1 Introduction

All biological systems are composed of a huge number of molecular interactions, and the goal of any drug is to influence a subset of these interactions associated with a particular disease. As such, computational drug discovery requires a deep understanding of all interactions - both the interaction between the drug and target molecules (efficacy) and the many more potential interactions with molecules that must be avoided (specificity).

Recent progress has been made in predicting the interactions between pairs of molecules. The problem, however, is that these advanced methods are difficult to scale, and the number of potential interactions between molecules in human biology is immense. Focusing on an important subset of these interactions - protein vs protein interactions (PPIs), it is estimated that there are *∼* 10^5^ different proteins in the human proteome, which means there are *∼* 10^10^ possible PPIs. Many recent innovations have increased the scale of experimental methods for detecting PPIs, such as yeast two-hybrid assays and co-immunoprecipitation, but the current state of the art is capable of measuring only *∼*10^5^ possible interactions. Computational approaches have also seen major advancements in recent years, particularly methods that utilize protein structure prediction algorithms such as AlphaFold2 [14] and RoseTTAFold [3]. While powerful and more scalable than experimental methods, these techniques rely on querying computationally intensive models for every possible protein-protein pair. As such, they have thus far failed to scale to the entire PPI network, with predicted networks built by only querying *∼* 10^6^ potential interactions [24, 26].

Vector representations offer an effective solution to the problem of inferring entire PPI networks. The interaction between two proteins is a complicated, non-linear process, but this does not preclude the existence of a high-dimensional vector space where this interaction can be approximated as a dot product (a quick and low-cost calculation). If all proteins in a proteome are mapped into such a vector space, then the entire PPI network can be inferred nearly instantly by calculating the dot product between all vector representations.

In this paper, we introduce Deep Kernel Inversion (DKI) - a novel deep learning model that projects each protein into a high-dimensional vector space using only the structure of the protein. In this vector space, the dot product between any two vectors is a highly accurate prediction of whether the vectors’ corresponding proteins will interact. Each protein in a proteome needs to be embedded into a vector space only once. Once embedded, predicting interactions between a protein and any other already-embedded proteins becomes a computationally trivial task of calculating the dot product between vectors. In other words, this changes the task of predicting an entire PPI network from *O*(*n*^2^) to *O*(*n*). For the human proteome, this means a *100,000-fold* reduction in computation, making it feasible for the first time to predict all possible human protein PPIs accurately. Additionally, this approach can be applied to interactions between all molecules, not just proteins, and it could potentially capture an even richer set of properties beyond static interactions. See Fig 1 for a conceptual description of how DKI predicts interactions using vector representations.

**Figure 1:**
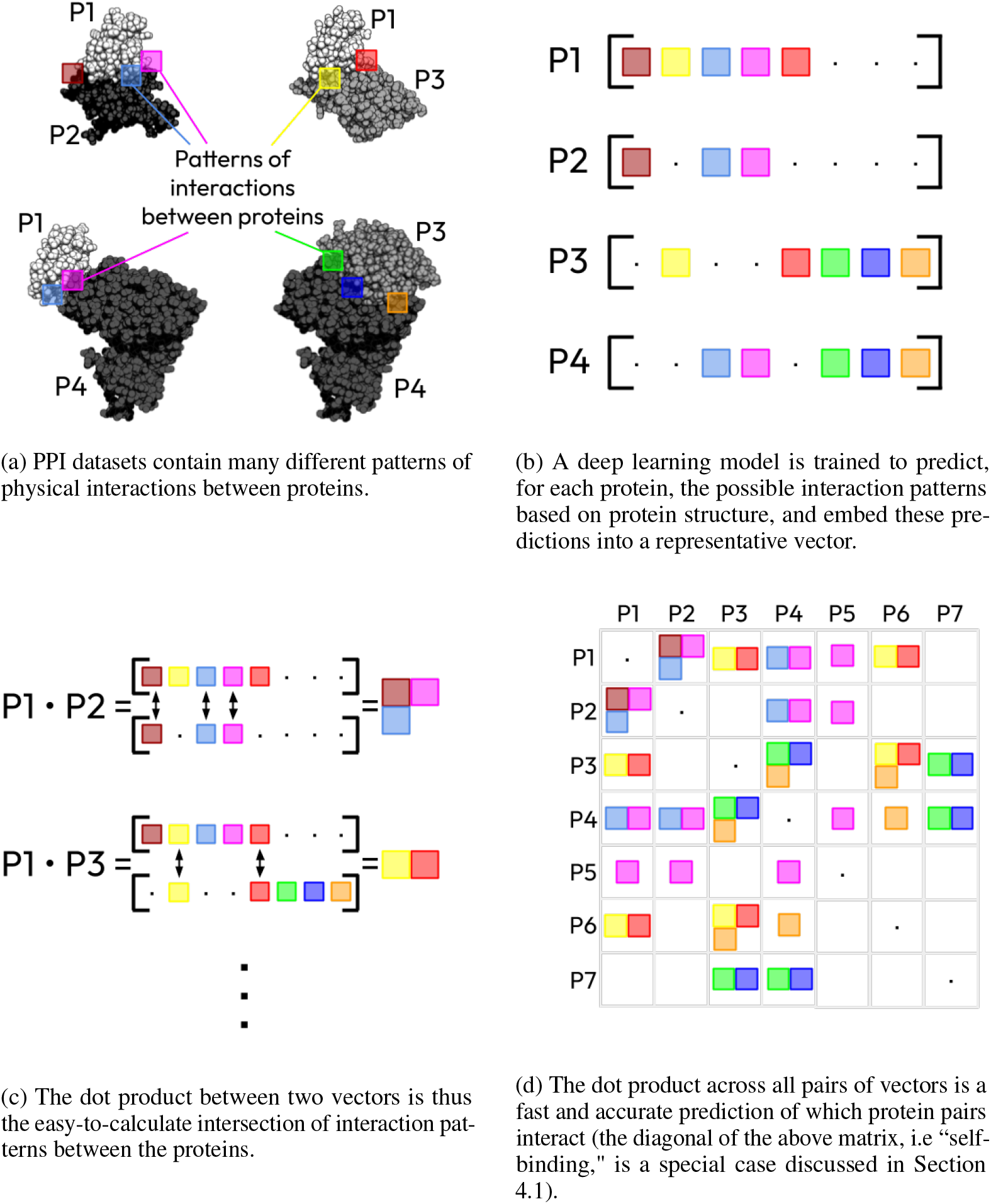
A conceptual description of how DKI uses vector representations to predict protein interactions. The goal of DKI is to encode all possible ways a protein could interact with another protein into a vector space so that a dot product between vectors is predictive of which interaction patterns two proteins have in common.

The key contributions of our work are as follows:

- **Novel approach to scaling physical interactions**

DKI represents a generalizable framework for scaling any number of physical interaction properties between large numbers of molecules of many different types.

- **Proteome-scale PPI network construction**

DKI allows for rapid and accurate prediction of entire PPI networks.

- **State-of-the-art accuracy**

We compare our approach to multiple past methods for PPI network prediction against several experimentally verified PPI networks, achieving best-in-class accuracy.

- **Accurate binding affinity prediction**

We show that DKI can be extended to predict not just bind/no-bind interactions between proteins but also the binding affinity of the interactions.

- **Full proteome PPI network**

We apply DKI to a large number of human proteins, mapping 147K interactions between 20K proteins.

Our results underscore the potential of deep learning-generated vector representations in revolutionizing computational molecular interaction prediction. The combination of high accuracy and computational efficiency opens new avenues for large-scale studies of molecular interaction networks, accelerating biological discovery and therapeutic development.

## 2 Background

There is a broad body of work related to predicting interactions between proteins. Many approaches use co-evolutionary data, essentially relying on the interactions between similar proteins in other species. In particular, multiple sequence alignment (MSA) is used to identify proteins with similar sequences or subsequences from which properties such as structure or interactions can be inferred. This inference has been done through statistical methods [7, 15, 17, 22, 28], traditional machine learning [4, 36, 37], and deep learning [8, 10, 13, 23, 29]. Given the success of deep learning-based methods for predicting protein structures, such as RoseTTAFold [3] and AlphaFold2 [14], many methods have been built directly on top of these tools with varying degrees of success [6, 9, 12, 16, 24, 26].

The challenge with MSA or co-evolutionary-based approaches is multifold. First, MSA is computationally expensive. This leads to some methods relying on a tiered system where quicker, less accurate models first process potential PPIs, and then the MSA-dependent models take a second pass over the highest-scoring candidates [12, 24]. Even with attempts to accelerate the computation, it remains the case that every protein pair within a proteome must be considered. This means querying a model billions of times, and none of these methods are capable of handling that scale.

Another challenge with MSA-dependent methods is that they are less effective with proteins that have fewer matching sequences, and especially with novel or unseen proteins. A major use case of PPI prediction at Nosis, for example, is in designing *de novo* proteins (novel proteins that are dissimilar to any protein found in nature). In particular, we use DKI to predict all interactions a newly designed protein will have with 120K different extracellular epitopes within the human proteome in order to optimize their specificity. With MSA-based methods, this would be impossible, both because no similar sequence to the designed protein has ever been observed, and because querying a model 120K times at each step in our iterative protein design process is unfeasible. In summary, MSA-based methods are simply extrapolating previously observed patterns, and they are prohibitively slow in doing so.

### Kernel-based methods in machine learning

Kernel functions are essentially similarity measures between items within a set, and they are used throughout machine learning [1, 11, 33]. Many machine learning algorithms work by making a series of comparisons between items within a dataset, but these comparisons are usually made via a dot product, which leads to models only being capable of finding linear patterns. It has been shown, however, that nonlinear kernels with certain properties

[19] can be substituted for these dot products, leading to more sophisticated pattern finding. The well-known “kernel trick” in Support Vector Machines is such an example.

The interchangeability of nonlinear kernels with dot products arises from the equivalence these kernels have to dot products in a much higher-dimensional vector space. What is a nonlinear comparison in lower dimensions is equivalent to a linear comparison when projected into a much higher dimensional space. We expand on this idea here. The method presented in this paper essentially inverts the kernel trick—starting with a nonlinear and expensive-to-calculate kernel (i.e., the likelihood that two proteins will interact), we find a higher-dimensional vector space in which the dot product is (near) equivalent to the kernel. In doing so, we only need to project each protein into this vector space once, and then simply by taking the dot product between all proteins, we can infer all interactions with no additional querying of our model.

Vector representations are also related to the idea of higher-dimensional encodings. Vector representations of words were heavily used in the early days of deep learning-based natural language processing because they could capture additional semantic information within the vectors themselves [20, 21, 25]. This meant that by building a vector space based on word co-occurrences across text, words could be fed into neural networks with more information than a simple one-hot encoding scheme. With word encodings, however, the enormity of data and the fact that words co-occur with many other words to varying degrees allowed the vectors to be inferred directly. There was no need to “predict” each word’s vector or map a word’s properties to its vector. Here, protein interactions are highly sparse, with most proteins having zero or one interaction that has been captured in structure. Thus, we must rely on deep learning to map the structure of the protein to its appropriate projection into the vector space.

## 3 Results

DKI offers significant improvements in computational efficiency over existing methods for PPI networks. These gains are accompanied by state-of-the-art performance in accuracy, as well as deeper insights into the nature of PPIs.

### 3.1 Performance benchmarks

We applied DKI to several validated PPI datasets and compared its performance to multiple recently published methods. All of these methods rely on MSA and, as such, exhibit variability in performance across datasets depending on the number of matched sequences available for the PPIs. Additionally, each method requires querying the model once for each protein pair rather than once for each protein, resulting in a much longer runtime. Additionally, our method demonstrates superior accuracy.

First, PEPPI [4] is a machine learning pipeline that utilizes multiple models based on structural and sequential similarities, which are then used as inputs into a naive Bayesian classifier. We applied this model and our DKI model to five different yeast PPI datasets available at the Yeast Interactome Project: (http://interactome.dfci.harvard.edu/S_cerevisiae/). These datasets had overlapping examples with our training dataset (see Section 4.3), so those examples were removed before running both models. Consistent with the methods in [4], we tested the models on a balanced dataset— for each true PPI, we randomly sampled one pair of non-interacting proteins as a negative example.

Figure 2 shows the performance of PEPPI versus DKI. Across every dataset, our model demonstrates superior performance. Additionally, DKI completed in minutes for all datasets using a small GPU, whereas PEPPI could only run on a traditional CPU and took multiple days to complete (with most of the CPU time spent on running MSA).

**Figure 2:**
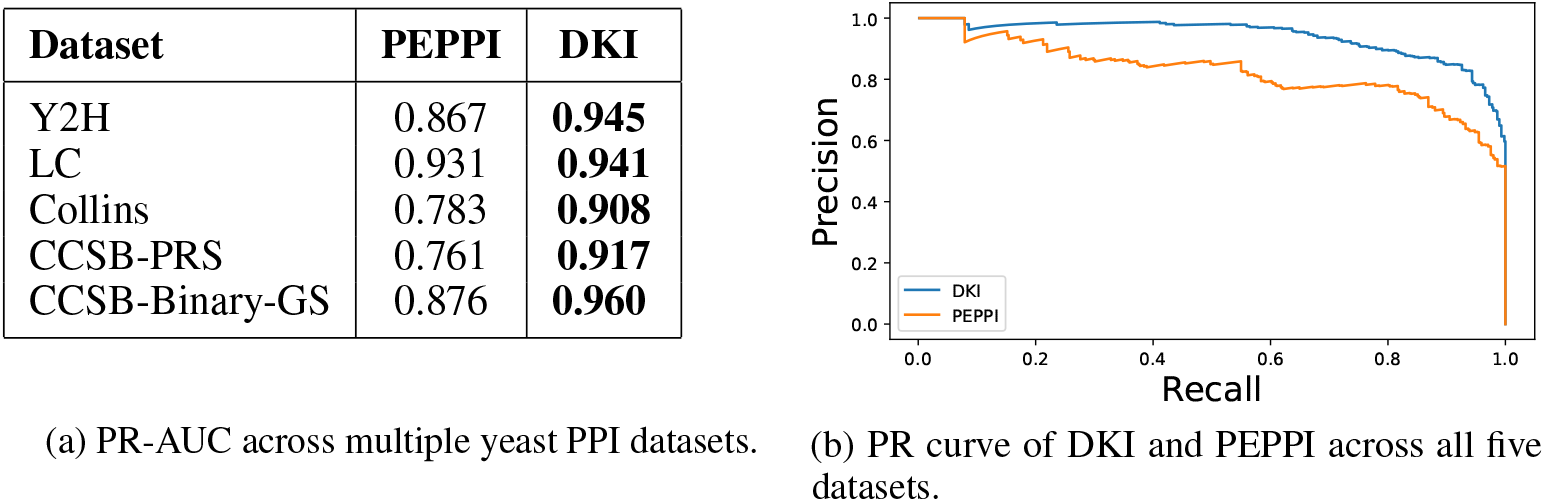
Performance (PR-AUC) of our Deep Kernel Inversion (DKI) model versus PEPPI [4] in predicting protein-protein interactions across multiple yeast interactome datasets.

Next, we compared DKI to the method described in [24], which utilizes AlphaFold2, RoseTTAFold, and coevolution of proteins across species (referred to as “Directed-Coupling Analysis” or DCA) to predict protein-protein interactions among human mitochondrial proteins [27]. We applied DKI to the same dataset to compare our performance with the results reported in their paper. As done in [24], we used a positive-to-negative example ratio of 1:100. The results are shown in Table 1. The DKI model outperforms the three different methods described in [24] by a wide margin. Additionally, [12] applied similar methods to the yeast proteome, noting that the most accurate AlphaFold2-based method would take up to 1 million “GPU hours” to predict the interactions among the 4K proteins in the yeast proteome. DKI takes less than one hour on a modestly powered GPU.

**Table 1:** Performance of DKI in predicting PPIs between human mitochondrial proteins compared to three different methods developed in [24]. The positive-to-negative example ratio used was 1:100.

| Model | PR-AUC |
| --- | --- |
| DKI | **0.304** |
| AF2 | 0.132 |
| RoseTTAFold | 0.113 |
| DCA | 0.014 |

In addition to these two works, several other papers have developed methods for predicting PPIs and reported performance on various datasets [6, 8, 12, 13, 23]. We have not compared our model head-to-head with all of these papers due to reasons such as datasets not being available, code not being accessible, and/or our inability to remove overlapping examples from their training data to prevent data leakage. However, in every case, the performance results reported are either inferior or not superior to what we have presented here. DKI represents a many-orders-of-magnitude speedup in the computation of PPI networks without sacrificing accuracy compared to the current state of the art.

#### Binding affinities

Not all protein-protein interactions are equal. Different interacting protein pairs have affinities (denoted as *K*_*d*_) that vary from 10^*−*3^ molar to 10^*−*15^ molar. The affinity between two interacting proteins is a function of how quickly, how often, and for how long the proteins remain bound to each other, and thus plays a crucial role in molecular dynamics.

We applied DKI to the problem of predicting binding affinities between interacting proteins by modifying the model to output a predicted log(*K*_*d*_) instead of a binary bind/no-bind probability. We used PDBBind (http://www.pdbbind.org.cn/index.php), a well-studied collection of many molecular interactions and their experimentally measured affinities, to first fine-tune and then test the model (ensuring that the test examples had minimal or no structural similarity to examples used for training and fine-tuning). The results can be seen in Figure 3. DKI achieved a Pearson correlation coefficient of *R* = 0.677.

**Figure 3:**
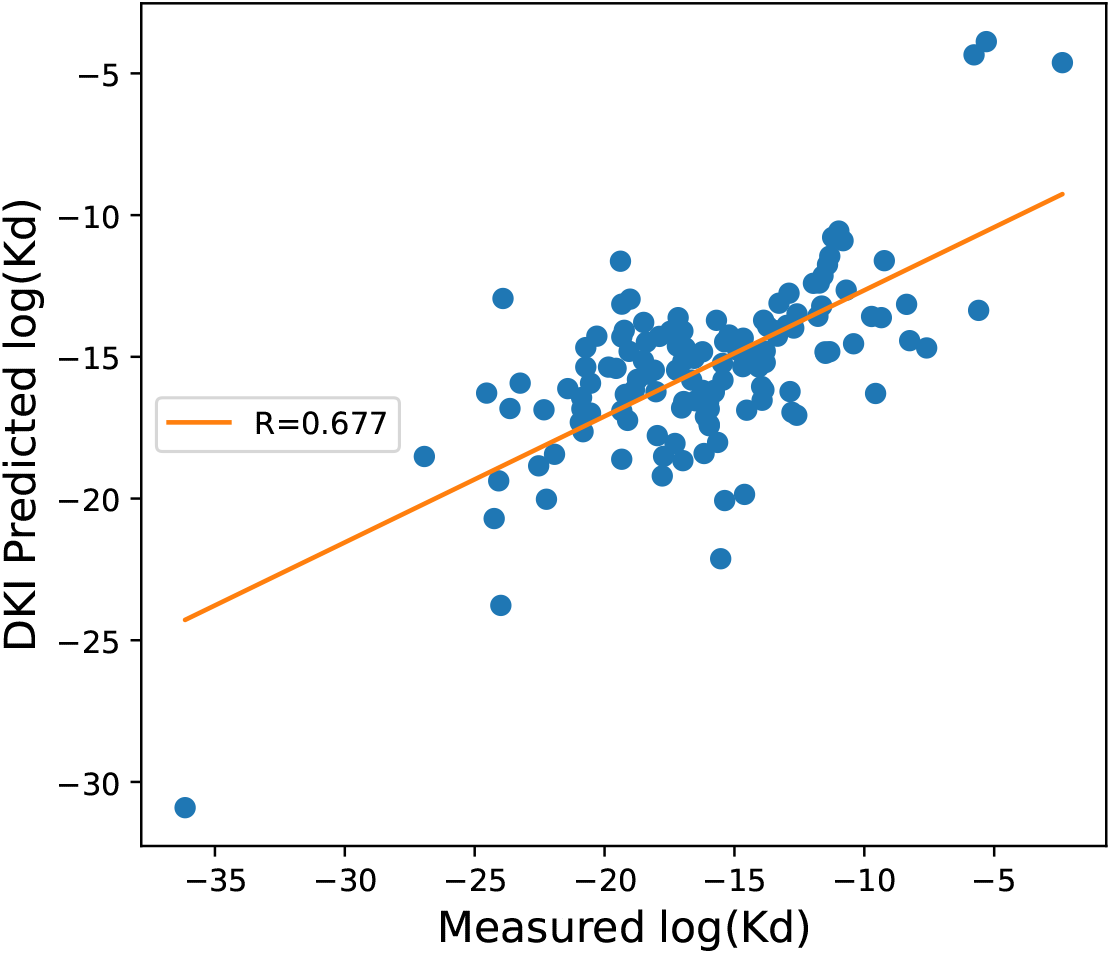
DKI predicted binding affinities versus measured affinities for protein-protein binding.

This result places DKI among the top two entrants in the CASF-16 binding affinity prediction competition [30], though since then, multiple groups have reported better performance (notably, Nosis also has an internal model for predicting protein binding affinity that also achieves superior performance). However, DKI’s ability to predict binding affinities for every single protein-protein interaction across an entire proteome is a significant advantage. With all other methods, which rely on querying the model with every protein pair, this would be infeasible.

### 3.2 Full-proteome PPI network construction

Lastly, to demonstrate the scalability of DKI, we predicted a complete PPI network for the entire set of human protein structures in the AlphaFold Protein Structure Database [34]. In total, this network includes 20,425 proteins, which implies there are 208.6 million possible interactions that could exist. According to [12], it would take somewhere between 90 million and 900 million GPU hours to apply AlphaFold2 to all of these possible interactions. In contrast, DKI ran on 4 Nvidia A100 GPUs for **4.5 hours** to complete the entire network.

In total, DKI predicted 147,291 interactions across these 20K structures. See Fig 4 for the distribution of the number of interactions per protein. We also used UMAP [18] to visualize the vector space of the proteins used to predict interactions, shown in Fig. 5. In this embedding, two proteins that are near each other are more likely to interact with the same other proteins but not necessarily to each other. In fact, it is not uncommon for adjacently embedded proteins to have a very low likelihood of interaction with each other. See Section 4.1 for a discussion of protein self-binding.

**Figure 4:**
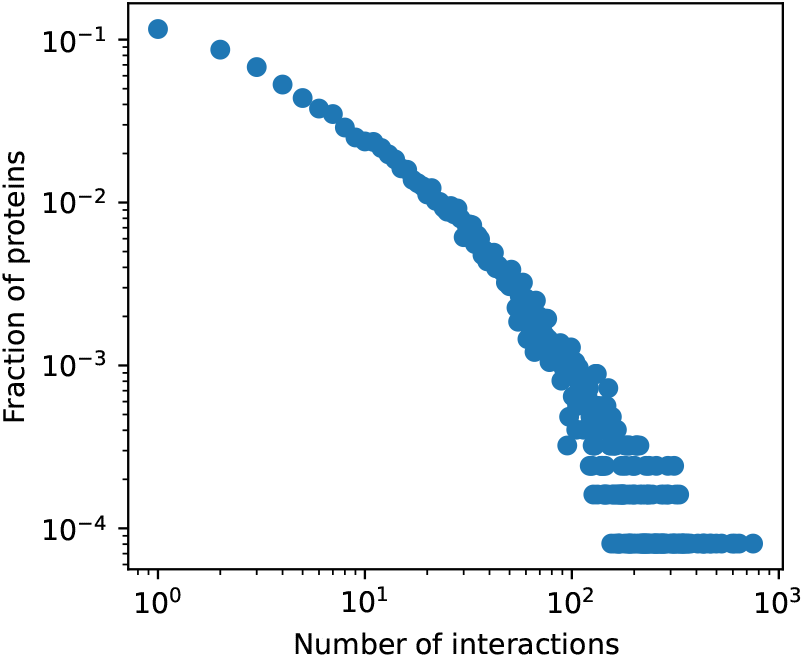
The distribution of the number of interactions predicted by DKI across proteins in the human Alphafold Protein Structure Database proteome. Roughly 12% of proteins have one predicted protein interaction.

**Figure 5:**
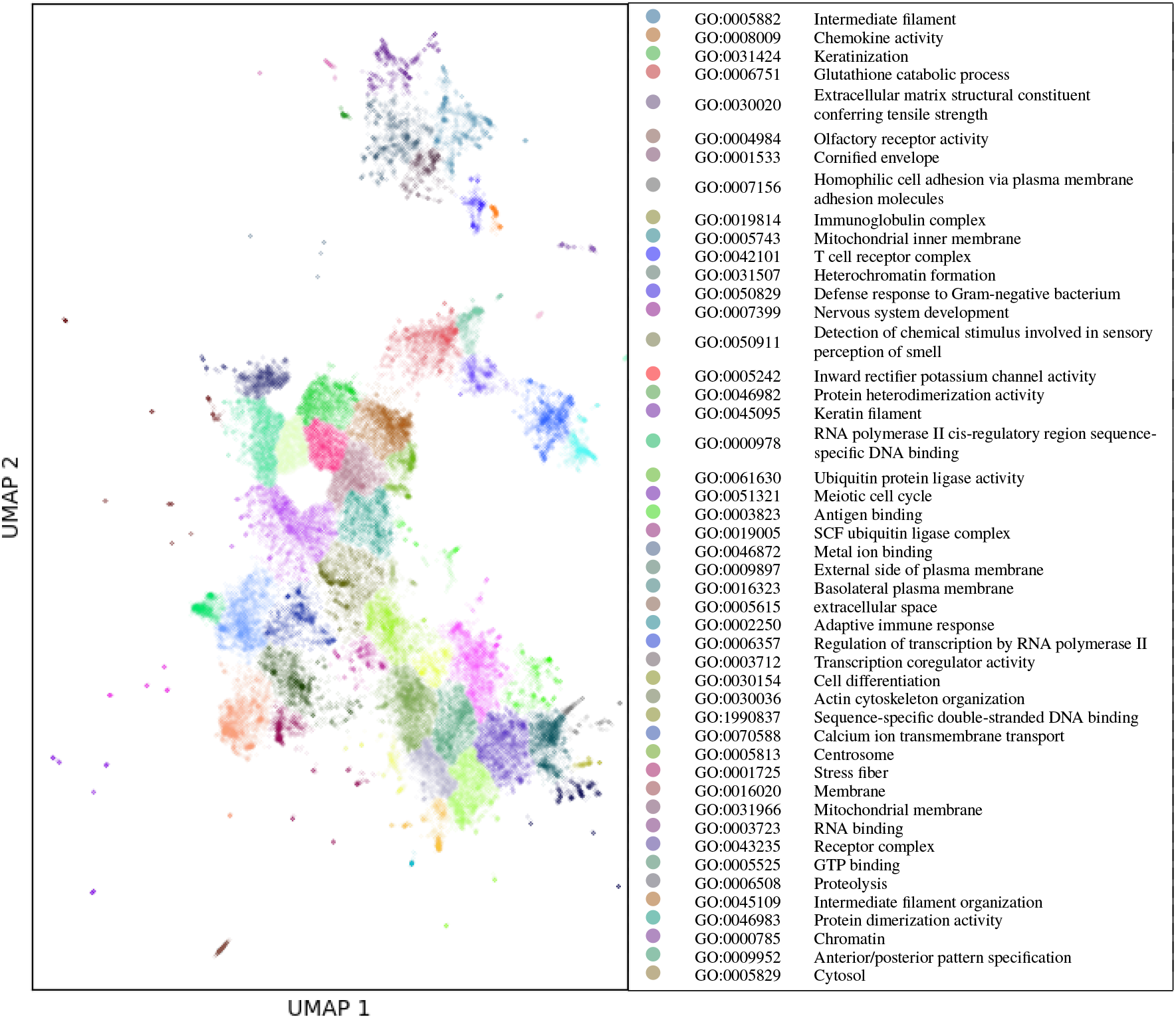
UMAP projection of the vector space embeddings for the human Alphafold Protein Structure Database. The clusters are labeled based on which Gene Ontology term [2] showed the most differentiated occurrence for proteins within a cluster compared to all other proteins.

To confirm the common binding of nearby proteins, we clustered the embeddings and labeled the resulting clusters based on which Gene Ontology term [2] showed the most differentiated occurrence for proteins within a cluster compared to all other proteins. Indeed, many of the differentiating Gene Ontology terms represent binding to proteins and other molecules, such as “Antigen binding”, “Sequence-specific double-stranded DNA binding”, and “Ubiquitin protein ligase activity”. In other words, many of the embeddings within these clusters are defined by binding to the same 3rd-party molecules.

This proximity of the embeddings based on common binding (but, again, this does not necessarily mean they bind to each other) makes DKI particularly useful in predicting the specificity of therapeutics. Designing therapeutics for specificity, or the degree to which a therapeutic molecule avoids interacting with off-target proteins) is a significantly harder problem than on-target binding affinity because it means optimizing interactions with more than 100,000 epitopes throughout human biology. This very challenge, in fact, is what originally motivated the invention of DKI at Nosis. We mapped every extracellular epitope in the human proteome and calculated their vector embeddings using DKI. Then, when designing a new molecule targeting a specific epitope, we use embeddings adjacent to the target epitopes’ embedding to identify the proteins most likely to have off-target binding. This, in combination with using DKI to rapidly predict interactions between our designed molecule and every protein in the human proteome, means we design for specificity by minimizing *every* possible off-target interaction. To our knowledge, this is a singular capability.

## 4 Methods

Here, we describe the construction of the DKI model. The goal is to build a vector space in which all proteins are represented as vectors, and the likelihood of two vectors physically interacting is proportional to the dot product between their corresponding vectors. Specifically, let 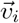 be the vector representing protein *i*, and let *b*_*i*_ be a bias scalar. Then

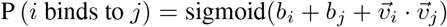

### 4.1 Self and anti-self binding

One immediate challenge in building such a vector representation of proteins is that the dot product of any vector with itself will be positive. Given that the vector space must be high-dimensional to capture all the variance and dynamics between interacting proteins, the vector’s self dot product will likely be higher than its dot product with most other proteins’ vectors. This leads to a prediction that all proteins are self-binding dimers, which is not true. Additionally, many protein pairs with similar or homologous structures may be incorrectly predicted to interact with high probability.

To address this, our representative vectors are complex-valued, i.e.,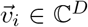for all *i*, where *D* is the vector space dimension. The imaginary components of the vector entries capture “anti-self” binding properties—such as convex/concave shapes, hydrogen-bond donating/accepting atoms, and other structural elements that interact strongly with their complement and not with themselves. The real components, on the other hand, capture self-interacting elements like hydrophobic surfaces and protein interfaces that match in surface area.

Additionally, as mentioned, the vector space needs to be high-dimensional to capture the nonlinear dynamics of protein-protein interactions. Therefore, we found it necessary to include a scaling factor in the dot product, as done in [35], for effective training of the model. This leads to the final expression:

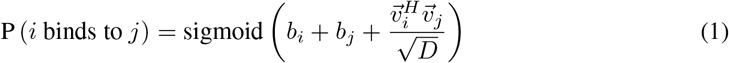

Where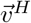is the conjugate transpose of vector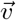, i.e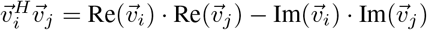. If, for example, a protein surface has a large number of positively charged residues arranged in a concave shape, then Re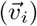 might be small in magnitude while Im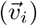 is large. When dotted with itself, the imaginary component would dominate, resulting in a low predicted probability of binding. In contrast, if the protein surface were flat and hydrophobic, the real component would likely dominate.

### 4.2 Mapping Structure to Vectors

Unlike other problems addressed using vector representations, PPI networks are highly sparse. Thus, each protein’s vector space projection cannot be inferred directly but must be predicted via a deep learning model, which we outline below.

#### Model inputs

The probability of two proteins binding when coming into contact should be determinable solely based on their respective structures. Our model depends exclusively on the primary amino acid sequence and backbone coordinates (i.e., the coordinates of the *C*_*α*_ atoms). We have developed a companion model that takes all atoms of the protein as input, but its design includes several innovations that we have chosen to keep as trade secrets. Due to our need for the model to work for *de novo* designed proteins and to be scalable to full proteome PPI networks, including multiple sequence alignment (MSA) or protein templates as input would be prohibitive.

Given the large variability in protein sizes, encoding the entire protein into a single vector might not be feasible. This can result in various surfaces on the protein that could interact with other proteins. In one version of the DKI model, we addressed this by including specific “binding site” residues of the protein (residues that are within 8 angstroms of the other protein upon binding) as model inputs. Since the binding site is often unknown for many different PPIs, we built a separate deep learning model to predict likely binding sites on a protein. Predicting the binding site on one protein without knowing the other protein is challenging. However, at Nosis, our primary focus is on designing small protein ligands (less than 100 residues) that bind to larger receptor proteins. This means the binding sites are of specific sizes, reducing variability and enabling our binding site predictor to achieve a per-residue PR-AUC > 0.95. Here, with the goal of modeling all PPI, we do not have the benefit of consistent binding site size, and an equivalent binding site classifier was only able to achieve a PR-AUC=0.79. This would mean that the binding site inputs into our general PPI model would be highly variable.

We found it more effective to minimize variability in the inputs and let the general model encode the variability in the resulting vector space. Thus, instead of including the entire binding site, we include a “seed residue” for the protein. The seed residue is a surface residue (within 1 angstrom from the protein surface) that is minimally distant from all other residues in the *other* protein. This is similar to “hot residues,” although we do not require the residue to be crucial for the interaction. It represents the “center” of where the binding site might be. In constructing a PPI network, we generate a vector encoding for every surface residue in each protein. The probability that two proteins interact is the maximum probability predicted across all of their respective seed residue encodings.

#### Model architecture

We developed a novel deep learning model to map a protein’s structure to its vector space projection in C^*D*^. First, the mapping from structure to vector must be translationally and rotationally equivariant. This is achieved by restricting the input features to the amino acid types and the Euclidean distances between the *C*_*α*_ backbone atoms of residues.

As in [3, 14], we use an amino acid pairwise encoding architecture. However, our architecture differs significantly from these works. First, while their goal is to predict protein structure, our model uses protein structure as an input. Second, both works rely heavily on MSA (using other known protein structures with similar amino acid sequences), whereas our model must work for *de novo* proteins (proteins dissimilar to any protein found in nature), so MSA is not used. Lastly, our neural network architecture differs substantially due to these other points of contrast.

Our architecture consists of two parts: the residue pair encoder and a self-attention-based “reducer.” The residue pair encoder learns vector representations for each pair of residues/amino acids in the protein structure. The pair encoder comprises 12 blocks of layers, each block being a transformer that performs two self-attentions—one with the first residue constant and the second varying, and a second with the second residue constant and the first varying. Specifically, in any given layer, the attention vector for residue pair *i, j* is

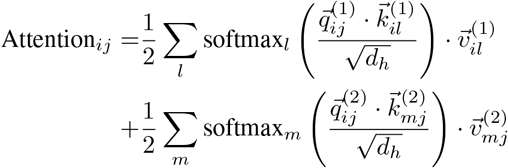

Where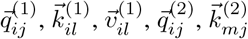and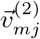are vectors of dimension *d*_*h*_. We use *d*_*h*_ = 128. Note that the above equation represents “single-headed” attention. We do use multi-headed attention, but the specific notation was omitted here for clarity. The rest of the transformer architecture is standard, including layer normalization, feed-forward networks, and skip connections.

The output of the neural network must be a single vector (i.e., the protein’s vector representation). Therefore, on top of the 12 residue pair transformers, we use an attention reducer layer to “reduce” the *n* ^2^ residue pair encodings into a single vector. This is also achieved using self-attention:

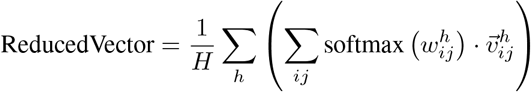

where *H* is the number of attention heads (we use *H* = 8), 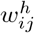is a scalar, and 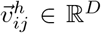, both generated by single-layer linear feed-forward networks using the residue pair *ij*’s encoding as inputs.

The ReducedVector is then passed through LayerNormalization and a two-layer feed-forward network. To generate a complex vector (as discussed above), we produce the real and imaginary components of the vector separately using independent two-layer feed-forward networks. See Fig 6 for an architectural diagram.

**Figure 6:**
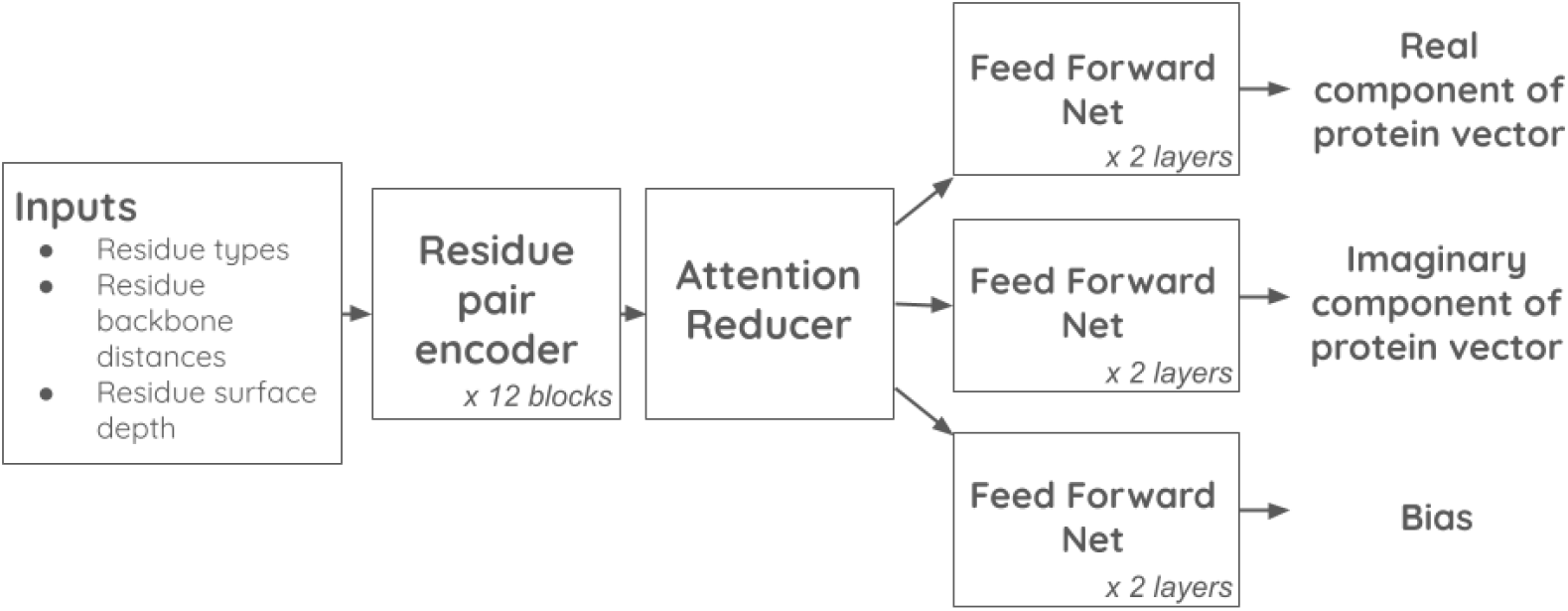
Diagram of the neural network used to map proteins into the interaction vector space.

### 4.3 Model Training

Projecting protein surfaces into a high-dimensional vector space enables significant improvements in PPI network inference and also introduces a more efficient training process. However, the high-dimensional vector space presents its own set of unique training challenges.

#### Loss function

Predicting protein-protein interactions is a binary classification problem—given two proteins, do they interact (yes or no)? A naive training process might randomly sample pairs of proteins, some of which are known to interact while others are negative examples (two proteins sampled randomly with no known interaction). The high sparsity of interactions means the false negative rate of this sampling is negligible. The forward-pass step in backpropagation must output predicted vectors for every protein in the sampled pairs. Once generated, each vector within a training batch can be compared to the other vector in its pair as well as to the vectors in other pairs with no additional computation. Again exploiting the sparsity of PPI networks, we can add these comparisons to the loss without increasing the compute requirements for backpropagation.

More specifically, given a training batch size of *b*, let *l*_*ij*_ *∈* {0, 1} indicate whether the *i*-th and *j*-th training examples in the batch are known to interact (i.e., positive training examples). The binary cross-entropy loss is then

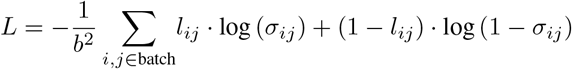

where

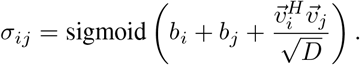

This expression can be computed entirely through matrix operations, so when running on GPUs, inter-vector comparisons add comparatively negligible time to the training step.

We used a batch size of 80 proteins during training, which means the traditional classifier approach to training would train on 80/2 = 40 protein *pair* examples per batch. With this new approach, each batch contains 80 *∗* 79/2 = 3160 protein pair examples, representing a significant gain in training efficiency. Since nearly all inter-pair comparisons are labeled as negative examples, we only sampled positive pairs to populate the batch. Thus, each sampled protein has one positively labeled comparison in the batch and 79 negative ones.

#### Training data

To train the vector encoder, we extracted a dataset of structural interactions between different proteins from publicly available PDBs. For each PDB, if two chains had three or more residues in contact (C_*α*_-C_*α*_ distance < 8 angstroms) and both chains were mapped to different UNIPROT IDs, we included the protein pair in our dataset. Each unique protein pair was included once, even if it appeared multiple times in the same or different PDBs. The result was 13,252 protein pairs. We partitioned the pairs such that 80% were in the training dataset, 10% were in a validation dataset (used for early stopping during training to prevent overfitting and for hyperparameter optimization), and 10% were used in a test set. We used Uniref90 clusters [31] to ensure that all pairs with sequential similarities were not split across datasets to prevent data leakage.

Initial training yielded poor results, with the model achieving a PR-AUC of 0.17 on the test set. The challenge lies in the required size of the vector space.

Since protein interactions can be complex and nonlinear, the vector space required to encode these interactions linearly must be large. Through use of the validation set, we determined that the optimal dimension for the vector space was *D* = 1024. However, larger vector spaces become more sparsely populated, and the Curse of Dimensionality [5] becomes a factor. In other words, a major disadvantage of DKI is that training is data-intensive. We tried to address this by loosening restrictions on including duplicates and chains not mapped to UNIPROT IDs in the dataset. This increased the dataset size to nearly 100K but decreased example quality, resulting in only a minor improvement in performance.

To address the massive data need, we used a proprietary dataset of structural protein-protein interactions developed by Nosis. The details about this dataset are considered trade secrets, but it includes 2.92 million relevant training examples of PPI. We pre-trained our model on this proprietary dataset (with special consideration to avoid data leakage) and then fine-tuned it on the 13K examples from public data. This approach significantly improved performance, achieving a PR-AUC of 0.57 (noting that the positive to negative example ratio is 1:79).

#### Predicting binding affinities

We modified the output of the DKI model and fine-tuned it on a protein binding affinity dataset. Specifically, the output was modified from Eq. 1 to

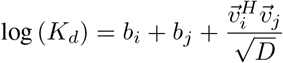

where *K*_*d*_ is the binding affinity of protein *i* to protein *j*. We used the subset of PDBBind protein-protein interactions where *K*_*d*_ was measured (as opposed to other measures of affinity), resulting in 1601 interactions. We split these interactions into training, validation, and test sets while ensuring Uniref90 partitioning was consistent with our other training datasets (see 4.3) to prevent data leakage. The binary PPI DKI model was fine-tuned to predict log(*K*_*d*_) using the training set. Figure 3 shows the results of the predictions against the test set. The fine-tuned DKI model achieved a Pearson correlation coefficient of *R* = 0.677.

#### Building the human proteome PPI network

We extracted from the Alphafold Protein Structure Database the 20,425 “V1” structures mapped to human UNIPROT records. In each structure we removed unfolded or low-confidence regions before embedding them into the vector space. We did this by removing any 4 or more consecutive residues with pLDDT<50.

As discussed in Section 4.2, each protein contains potentially multiple epitopes, or surfaces which could physically interact with another protein. An interaction between two proteins was predicted based on the max score across each proteins’ seed residue projections. In total, 57,120 seed residues were predicted to be interacting, from which 147,291 total protein-protein interactions were predicted.

Fig 5 is a UMAP projection of the 57K seed residue vectors (the real and imaginary components of each vector were put into separate vectors and then concatenated to form one vector of twice the dimension). Clustering was performed using using Leiden clustering [32]. Gene Ontology labels were identified by first calculating the percent of occurrence of each GO term across the unique proteins found within a cluster and then ranking the terms based on the log2 fold change of the occurrence rate in-cluster versus out-of-cluster. The clusters in the figure are ranked in descending order of the max log2FC, with the top cluster labeled “Intermediate filament” scoring a log2FC greater than 4.5.

## 5 Conclusion

Deep Kernel Inversion (DKI) is a novel approach to scaling molecular interaction predictions, offering speed improvements of several orders of magnitude over previous methods. What was once an *O*(*n*^2^) problem is now reduced to *O*(*n*). Additionally, this gain in speed does not come at the cost of accuracy, as DKI achieves state-of-the-art performance across multiple datasets.

Beyond speed, DKI is based solely on the structure of interacting proteins and does not rely on MSA or any other coevolutionary signals. This makes it particularly well-suited for characterizing interactions involving novel targets, binding sites, and novel compounds.

As demonstrated by its success in predicting drug properties like binding affinity, DKI can be used to design drugs with optimized affinity, specificity, and other properties required for therapeutic development.

## References

[1] Aizerman. Theoretical foundations of the potential function method in pattern recognition learning. Automation and remote control, 25:821–837, 1964.

[2] S. A. Aleksander, J. Balhoff, S. Carbon, J. M. Cherry, H. J. Drabkin, D. Ebert, M. Feuermann, P. Gaudet, N. L. Harris, et al. The gene ontology knowledgebase in 2023. Genetics, 224(1):iyad031, 2023.

[3] M. Baek, F. DiMaio, I. Anishchenko, J. Dauparas, S. Ovchinnikov, G. R. Lee, J. Wang, Q. Cong, L. N. Kinch, R. D. Schaeffer, et al. Accurate prediction of protein structures and interactions using a three-track neural network. Science, 373(6557):871–876, 2021.

[4] E. W. Bell, J. H. Schwartz, P. L. Freddolino, and Y. Zhang. Peppi: whole-proteome protein-protein interaction prediction through structure and sequence similarity, functional association, and machine learning. Journal of molecular biology, 434(11):167530, 2022.

[5] R. Bellman, R. Corporation, and K. M. R. Collection. Dynamic Programming. Rand Corporation research study. Princeton University Press, 1957.

[6] P. Bryant, G. Pozzati, and A. Elofsson. Improved prediction of protein-protein interactions using alphafold2. Nature communications, 13(1):1265, 2022.

[7] Q. Cong, I. Anishchenko, S. Ovchinnikov, and D. Baker. Protein interaction networks revealed by proteome coevolution. Science, 365(6449):185–189, 2019.

[8] G. Czibula, A.-I. Albu, M. I. Bocicor, and C. Chira. Autoppi: An ensemble of deep autoencoders for protein–protein interaction prediction. Entropy, 23(6):643, 2021.

[9] M. Gao, D. Nakajima An, J. M. Parks, and J. Skolnick. Af2complex predicts direct physical interactions in multimeric proteins with deep learning. Nature communications, 13(1):1744, 2022.

[10] L. Hallee and J. P. Gleghorn. Protein-protein interaction prediction is achievable with large language models. bioRxiv, pages 2023–06, 2023.

[11] T. Hofmann, B. Schölkopf, and A. J. Smola. Kernel methods in machine learning. 2008.

[12] I.R. Humphreys, J. Pei, M. Baek, A. Krishnakumar, I. Anishchenko, S. Ovchinnikov, J. Zhang, T. J. Ness, S. Banjade, S. R. Bagde, et al. Computed structures of core eukaryotic protein complexes. Science, 374(6573):eabm4805, 2021.

[13] K. Jha, S. Saha, and H. Singh. Prediction of protein–protein interaction using graph neural networks. Scientific Reports, 12(1):8360, 2022.

[14] J. Jumper, R. Evans, A. Pritzel, T. Green, M. Figurnov, O. Ronneberger, K. Tunyasuvunakool, R. Bates, A. Žídek, A. Potapenko, et al. Highly accurate protein structure prediction with alphafold. nature, 596(7873):583–589, 2021.

[15] H. Kamisetty, S. Ovchinnikov, and D. Baker. Assessing the utility of coevolution-based residue– residue contact predictions in a sequence-and structure-rich era. Proceedings of the National Academy of Sciences, 110(39):15674–15679, 2013.

[16] J. Ko and J. Lee. Can alphafold2 predict protein-peptide complex structures accurately? BioRxiv, pages 2021–07, 2021.

[17] D. S. Marks, L. J. Colwell, R. Sheridan, T. A. Hopf, A. Pagnani, R. Zecchina, and C. Sander. Protein 3d structure computed from evolutionary sequence variation. PloS one, 6(12):e28766, 2011.

[18] L. McInnes, J. Healy, and J. Melville. Umap: Uniform manifold approximation and projection for dimension reduction. arXiv preprint arXiv:1802.03426, 2018.

[19] J. Mercer. Functions of positive and negative type, and their connection the theory of integral equations. Philosophical transactions of the royal society of London. Series A, containing papers of a mathematical or physical character, 209(441-458):415–446, 1909.

[20] T. Mikolov. Efficient estimation of word representations in vector space. arXiv preprint arXiv:1301.3781, 2013.

[21] T. Mikolov, K. Chen, G. S. Corrado, and J. A. Dean. Computing numeric representations of words in a high-dimensional space, May 19 2015. US Patent 9,037,464.

[22] F. Morcos, A. Pagnani, B. Lunt, A. Bertolino, D. S. Marks, C. Sander, R. Zecchina, J. N. Onuchic, T. Hwa, and M. Weigt. Direct-coupling analysis of residue coevolution captures native contacts across many protein families. Proceedings of the National Academy of Sciences, 108(49):E1293–E1301, 2011.

[23] J. Pan, Z.-H. You, L.-P. Li, W.-Z. Huang, J.-X. Guo, C.-Q. Yu, L.-P. Wang, and Z.-Y. Zhao. Dwppi: a deep learning approach for predicting protein–protein interactions in plants based on multi-source information with a large-scale biological network. Frontiers in Bioengineering and Biotechnology, 10:807522, 2022.

[24] J. Pei, J. Zhang, and Q. Cong. Human mitochondrial protein complexes revealed by large-scale coevolution analysis and deep learning-based structure modeling. Bioinformatics, 38(18):4301–4311, 2022.

[25] J. Pennington, R. Socher, and C. D. Manning. Glove: Global vectors for word representation. In Proceedings of the 2014 conference on empirical methods in natural language processing (EMNLP), pages 1532–1543, 2014.

[26] D. Petrey, H. Zhao, S. J. Trudeau, D. Murray, and B. Honig. Preppi: a structure informed proteome-wide database of protein–protein interactions. Journal of molecular biology, 435(14):168052, 2023.

[27] S. Rath, R. Sharma, R. Gupta, T. Ast, C. Chan, T. J. Durham, R. P. Goodman, Z. Grabarek, M. E. Haas, W. H. W. H. Hung, P. R. Joshi, A. A. Jourdain, S. H. Kim, A. V. Kotrys, S. S. Lam, J. G. McCoy, J. D. Meisel, M. Miranda, A. Panda, A. Patgiri, R. Rogers, S. Sadre, H. Shah, O. S. Skinner, T.-L. To, M. A. Walker, H. Wang, P. S. Ward, J. Wengrod, C.-C. Yuan, S. E. Calvo, and V. K. Mootha. Mitocarta3.0: an updated mitochondrial proteome now with sub-organelle localization and pathway annotations. Nucleic acids research, 49(D1):D1541–D1547, 2021.

[28] S. Seemayer, M. Gruber, and J. Söding. Ccmpred—fast and precise prediction of protein residue–residue contacts from correlated mutations. Bioinformatics, 30(21):3128–3130, 2014.

[29] Song, X. Luo, X. Luo, Y. Liu, Z. Niu, and X. Zeng. Learning spatial structures of proteins improves protein–protein interaction prediction. Briefings in bioinformatics, 23(2):bbab558, 2022.

[30] M. Su, Q. Yang, Y. Du, G. Feng, Z. Liu, Y. Li, and R. Wang. Comparative assessment of scoring functions: the casf-2016 update. Journal of chemical information and modeling, 59(2):895–913, 2018.

[31] E. Suzek, Y. Wang, H. Huang, P. B. McGarvey, C. H. Wu, and U. Consortium. Uniref clusters: a comprehensive and scalable alternative for improving sequence similarity searches. Bioinformatics, 31(6):926–932, 2015.

[32] V. A. Traag, L. Waltman, and N. J. Van Eck. From louvain to leiden: guaranteeing well-connected communities. Scientific reports, 9(1):1–12, 2019.

[33] V. Vapnik. Support-vector networks. Machine learning, 20:273–297, 1995.

[34] M. Varadi, D. Bertoni, P. Magana, U. Paramval, I. Pidruchna, M. Radhakrishnan, M. Tsenkov, S. Nair, M. Mirdita, J. Yeo, O. Kovalevskiy, K. Tunyasuvunakool, A. Laydon, A. Žídek, H. Tomlinson, D. Hariharan, J. Abrahamson, T. Green, J. Jumper, E. Birney, M. Steinegger, D. Hassabis, and S. Velankar. Alphafold protein structure database in 2024: providing structure coverage for over 214 million protein sequences. Nucleic acids research, 52(D1):D368–D375, 2024.

[35] A. Vaswani, N. Shazeer, N. Parmar, J. Uszkoreit, L. Jones, A. N. Gomez, L. Kaiser, and I. Polosukhin. Attention is all you need. CoRR, abs/1706.03762, 2017.

[36] L. Wong, Z.-H. You, S. Li, Y.-A. Huang, and G. Liu. Detection of protein-protein interactions from amino acid sequences using a rotation forest model with a novel pr-lpq descriptor. In Advanced Intelligent Computing Theories and Applications: 11th International Conference, ICIC 2015, Fuzhou, China, August 20-23, 2015. Proceedings, Part III 11, pages 713–720. Springer, 2015.

[37] Z.-H. You, L. Zhu, C.-H. Zheng, H.-J. Yu, S.-P. Deng, and Z. Ji. Prediction of protein-protein interactions from amino acid sequences using a novel multi-scale continuous and discontinuous feature set. In BMC bioinformatics, volume 15, pages 1–9. Springer, 2014.

